# The emerging role of receptor trafficking in signalosome formation and sustained long-term Wnt/*β*-catenin signaling

**DOI:** 10.1101/2025.11.20.689481

**Authors:** Fiete Haack, Kevin Burrage, Adelinde M. Uhrmacher

## Abstract

Despite its central role in development, cell homeostasis, and cancer, the mechanisms that govern canonical Wnt signaling, particularly how receptor internalization and membrane organization shape signalosome formation and downstream pathway activation, remain a topic of ongoing debate. To resolve this longstanding controversy, we developed a mechanistic model that integrates seemingly contradictory experimental observations into a coherent explanatory framework. Using an optimized multi-level rule-based modeling approach the model explicitly captures lateral membrane organization, receptor complex dynamics, endocytic routing and downstream signaling. Simulations reveal a previously unrecognized compensatory mechanism in which rapid recycling of partially immobilized receptor assemblies amplifies and stabilizes signalosome formation. The results further indicate that internalization and recycling are essential for stable long-term ***β***-catenin activation, offering an explanation why these processes are required for excessive signaling in cancer cells. The model further predicts that ordered membrane domains enhance signaling not merely by colocalizing receptors and DVL but by increasing effective residence time through rebinding-driven cluster stabilization. Integrating diverse experimental measurements into a unified simulation framework quantitatively reconciles conflicting experimental reports on the necessity of receptor internalization for pathway activation. By resolving these debates and providing falsifiable predictions for how trafficking and membrane organization jointly tune pathway output, the study offers a mechanistic foundation for targeted modulation of Wnt signaling in developmental and disease contexts.

## 1 Introduction

The canonical Wnt pathway is an evolutionarily conserved cell-cell signaling cascade that crucially controls proliferation, migration, and cell-fate decisions in development and adult tissue homeostasis [1–3]. In its aberrant form, canonical Wnt signaling is critically involved in the pathogenesis of various diseases, such as neurodegenerative disorders, metabolic syndromes, autoimmune diseases, and cancer [4, 5].

In canonical Wnt signaling, the central regulatory step is the stabilization and nuclear translocation of the transcriptional co-activator *β*-catenin (Catenin *β*-1). Under basal conditions, i.e., when Wnt ligands are absent, *β*-catenin is continuously targeted for degradation by the destruction complex. This multiprotein complex includes the scaffold components Adenomatous Polyposis Coli (APC) and Axis Inhibition Proteins 1 and 2 (AXIN1/2), which recruit *β*-catenin, as well as the serine/threonine kinases Casein Kinase I *α* (CK1*α*) and Glycogen Synthase Kinase 3*β* (GSK3*β*) [6, 7]. These kinases sequentially phosphorylate *β*-catenin, which induces its ubiquitination and subsequent proteasomal degradation [8, 9]. Canonical Wnt signaling is initiated when Wnt ligands bind to their cell-surface target receptors Frizzled (FZD) and low-density lipoprotein receptor-related protein 5 and 6 (LRP). Ligand receptor binding induces the recruitment of Dishevelled (DVL) to the membrane, particularely to membrane microdomains enriched in cholesterol, phosphatidylinositol 4,5-bisphosphate (PI(4,5)P2) and caveolin-1 (so-called caveolae) [10–12]. These membrane regions promote signalosome assembly — a large complex, favoring LRP6 phosphorylation at multiple PPPSP motifs [13–16]. LRP6 phosphorylation creates docking sites for Axin, displacing it from the cytoplasmic destruction complex, hence inducing stabilization and nuclear translocation of the target protein *β*-catenin to regulate target gene transcription as part of a larger transcriptional complex [14, 17, 18]. Recent studies and comprehensive reviews emphasize that membrane trafficking and receptor dynamics are not passive background processes, but central regulators of Wnt signal strength, duration, and spatial patterning [19–21]. A major axis of regulation is receptor and ligand internalization by multiple endocytic routes (clathrin-mediated, caveolin-dependent, macropinocytosis, and other clathrin-independent processes) [22–24]. Whether endocytosis potentiates or attenuates signaling appears to be context– and cargo-dependent, and is still under discussion [25–27]. Both positive and negative roles have been reported: Ubiquitin ligases (e.g., ZNRF3/RNF43) and RSPO proteins induce FRZ/LRP6 endocytosis and membrane clearance through clathrin-dependent internalization, linking signal amplitude to trafficking by controlling cell surface receptor abundance [28–31]. On the contrary Wnt/FRZ/LRP6 signalosome is internalized through a caveolin-dependent (caveolae-mediated) endocytic route that is known to promote signal transduction: Several studies demonstrate, that disruption of Caveolin-1 or caveolae formation blocks LRP6 internalization and downstream *β*-catenin activation, underscoring that caveolar uptake is essential for sustained canonical Wnt signaling (eg. [32–35]). However, even though the vital role of internalization on Wnt signaling has been manifested by various studies over the last decade, the impact of receptor trafficking and, in particular, receptor recycling has been mostly neglected so far and remains elusive for Wnt signaling [27]. Numerous studies suggest that receptor trafficking plays a beneficial role in various signaling pathways, where continuous rounds of endocytosis and recycling protect receptors from degradation (e.g., [36–39]). Further receptor trafficking may amplify signal transduction by concentrating ligands and downstream adaptors in endosomes, which prolongs kinase activity (e.g., EGFR, *β*-adrenergic receptors), while endosomal transport can induce receptor sorting, based on distinct lipid environments [40–43]. Further, some experimental studies already imply a more central role of receptor trafficking in canonical Wnt signaling than previously anticipated [33, 44]. To explore whether receptor trafficking might also play a beneficial role in Wnt signaling, we developed a mechanistic computational model that combines the essential regulatory steps at the membrane with receptor trafficking dynamics.

Due to its central role in numerous biological processes and cellular regulations, as well as in its aberrant form, in cancer development, the Wnt signalling pathway has been the subject of various simulation models [45]. However, membrane dynamics have only been considered in a few models so far [46–50]; none of them comprises an extensive, pathway-specific representation of receptor trafficking in Wnt signaling. Modeling the receptor life cycle, including receptor/protein complex formation, ligand – or raft-induced internalization, recycling, and intracellular sorting, poses a major challenge for computational representations of canonical Wnt signaling. These processes, together with the resulting spatial heterogeneity in receptors, kinases, and other signaling components, critically shape the pathway dynamics. To capture these processes, we employed a compartmental modeling approach using ML-Rules, a multilevel rule-based language [51, 52]. Cellular and subcellular structures—such as the plasma membrane, raft domains, endosomes, and lysosomes—are represented as nested compartments with variable molecular content and attributes (e.g., radius, volume). ML-Rules enables the dynamic creation, merging, modification, and removal of compartments during runtime, allowing for a straightforward representation of processes such as endocytosis, recycling, and lysosomal degradation through declarative reaction rules [53]. Reaction kinetics may depend on compartment attributes, enabling volume-dependent rate kinetics and dynamic redistribution of molecular species. This approach provided us with the means to mechanistically incorporate the essential steps of the LRP6 receptor life cycle into a coherent, quantitative model.

To construct and calibrate the model, numerous experimental studies have been considered. Qualitative insights have been used to construct and structurally shape the model, while quantitative, time-dependent data were used for calibration and validation experiments, wherever available. The model was fitted concurrently to data regarding receptor complex dimerization, LRP6 phosphorylation, LRP6 internalization, and *β*-catenin aggregation. Integrating diverse experimental data from varying cell types, tissues, experimental conditions, and perturbations into a simulation model yielded comprehensive mechanistic insights into the role and impact of receptor trafficking.

Our quantitative modeling and simulation approach is designed to complement traditional experimental studies and conceptual reviews of Wnt pathway dynamics [5, 21]. Existing reviews synthesize extensive experimental findings into coherent but primarily qualitative conceptual models and typically offer little quantitative analysis. By contrast, simulation studies – such as [49, 54, 55] – are well suited to advancing quantitative understanding. Here, we use a simulation framework as a focal point to integrate diverse knowledge sources into a qualitatively and quantitatively coherent picture. This enables systematic exploration of existing hypotheses and provides a more coherent and comprehensive understanding of the membrane trafficking and receptor dynamics underlying canonical Wnt signaling.

## 2 Results

Here, we discuss aspects of receptor mobility and its impact on activation of the pathway, gained from our quantitative modeling. The mechanistic model of canonical Wnt signaling presented here was built using a bottom-up approach. Starting with a very simple model comprising a minimal set of reaction rules, we extended the model step-by-step by incorporating mechanisms reported in the literature. Our goal was to find a model for the initial steps in canonical Wnt signaling, i.e., Wnt binding, receptor aggregation, and signalosome formation, with parameter values within biologically reasonable limits that reproduce experimental measurements of the following processes at the same time (see Methods section): 1) receptor dimerization (LRP6 and FZ) after Wnt binding, 2) LRP6 phosphorylation, 3) LRP6 internalization, 4) lipid rafts size distribution and 5) *β*-catenin aggregation.

### 2.1 Quantitative model of canonical WNT signaling

In the following, we describe the model that resulted from the progressive development of the model. Note that we discuss only the main molecular interactions, the major dynamics, and key assumptions to facilitate access and understanding of the model. Individual mechanisms are depicted in Figure 1, where each arrow represents reaction rules implemented in the model together with the corresponding reaction rate constants [53]. The parameter values for the reaction rate constants are listed in Table 2.

**Fig. 1:**
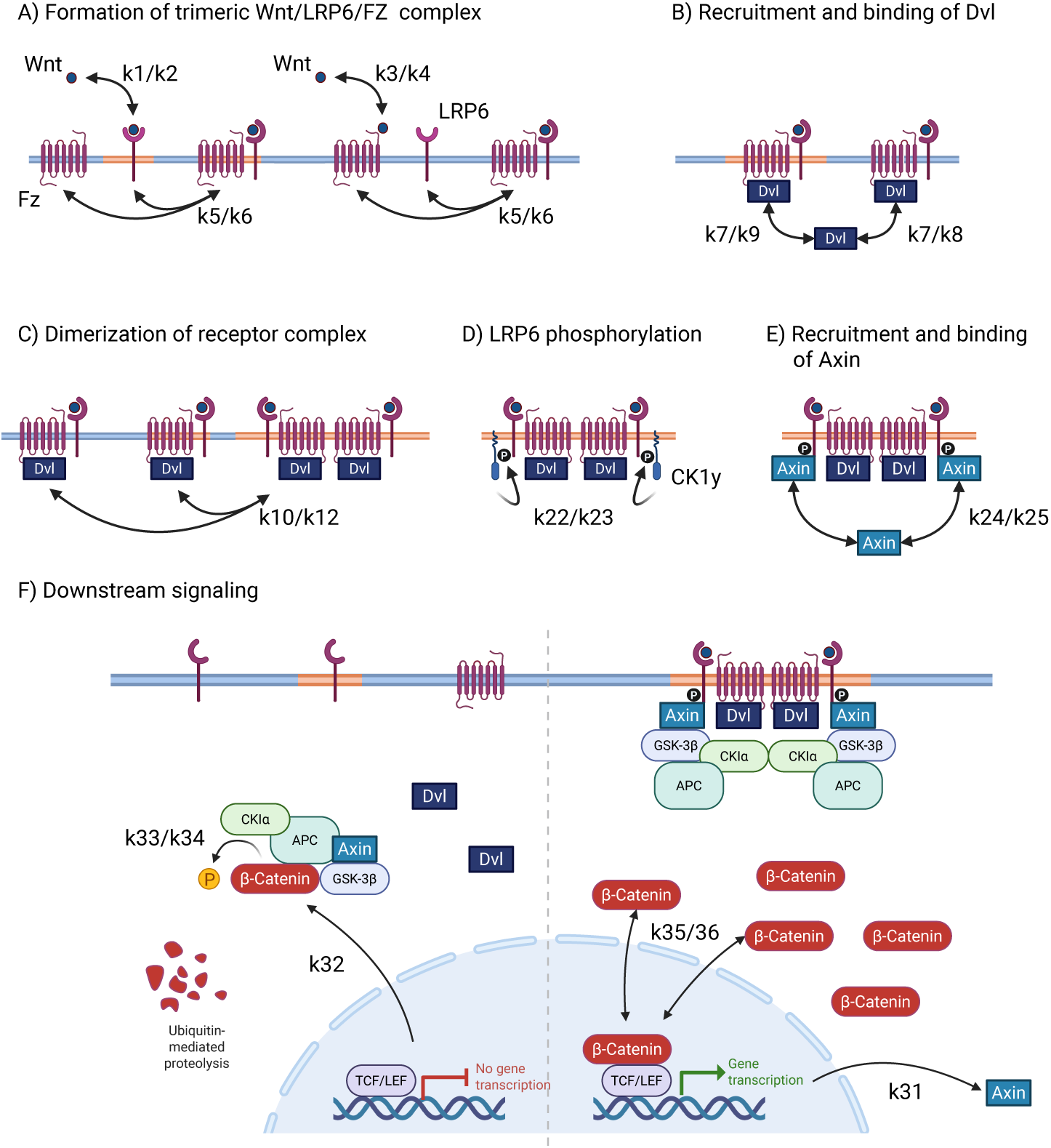
Simplified view of initial steps of signalosome formation and receptor activation after Wnt stimulus, as implemented in the model. The lower image (F) shows downstream signaling after successful signalosome formation in canonical Wnt signaling. The left side shows the inactive state (no Wnt stimulus), and the right side shows the activated state (with Wnt stimulus). Variable names of kinetic rate constants are associated to each arrow representing rules implemented in the model. Created in BioRender. Haack, F. (2025) https://BioRender.com/nyn65rn

**Table 1:**
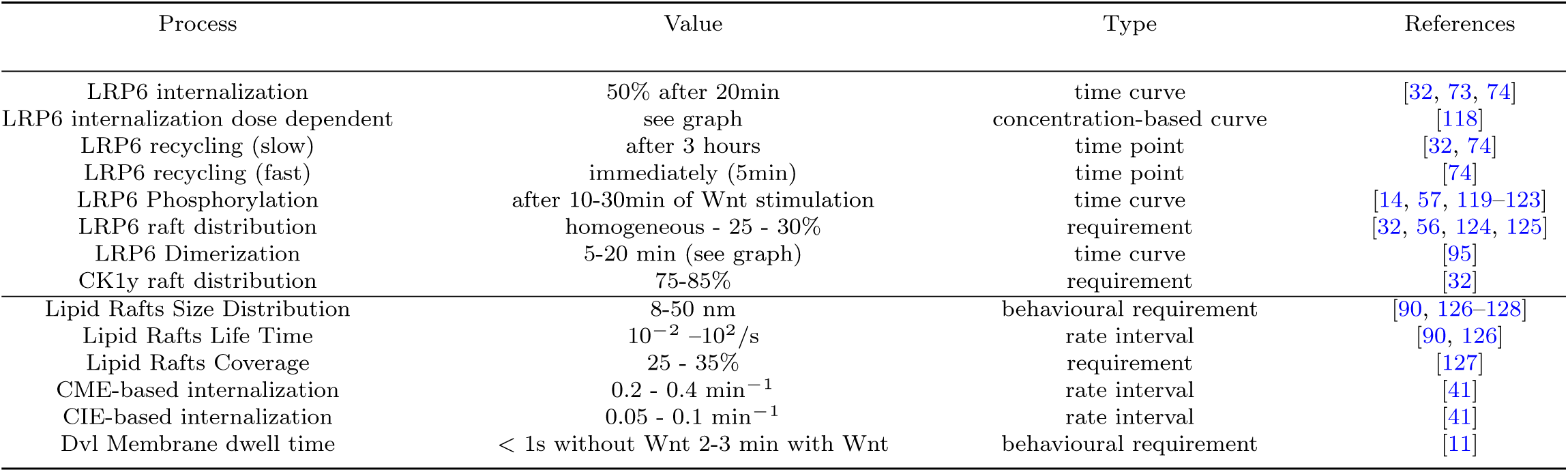
Table describing some model requirements, that were used during model development, based on experimental measurements. Requirements were, for instance, used as reference points or parameter value boundaries during model calibration, or for face validation of certain model dynamics.

**Table 2:**
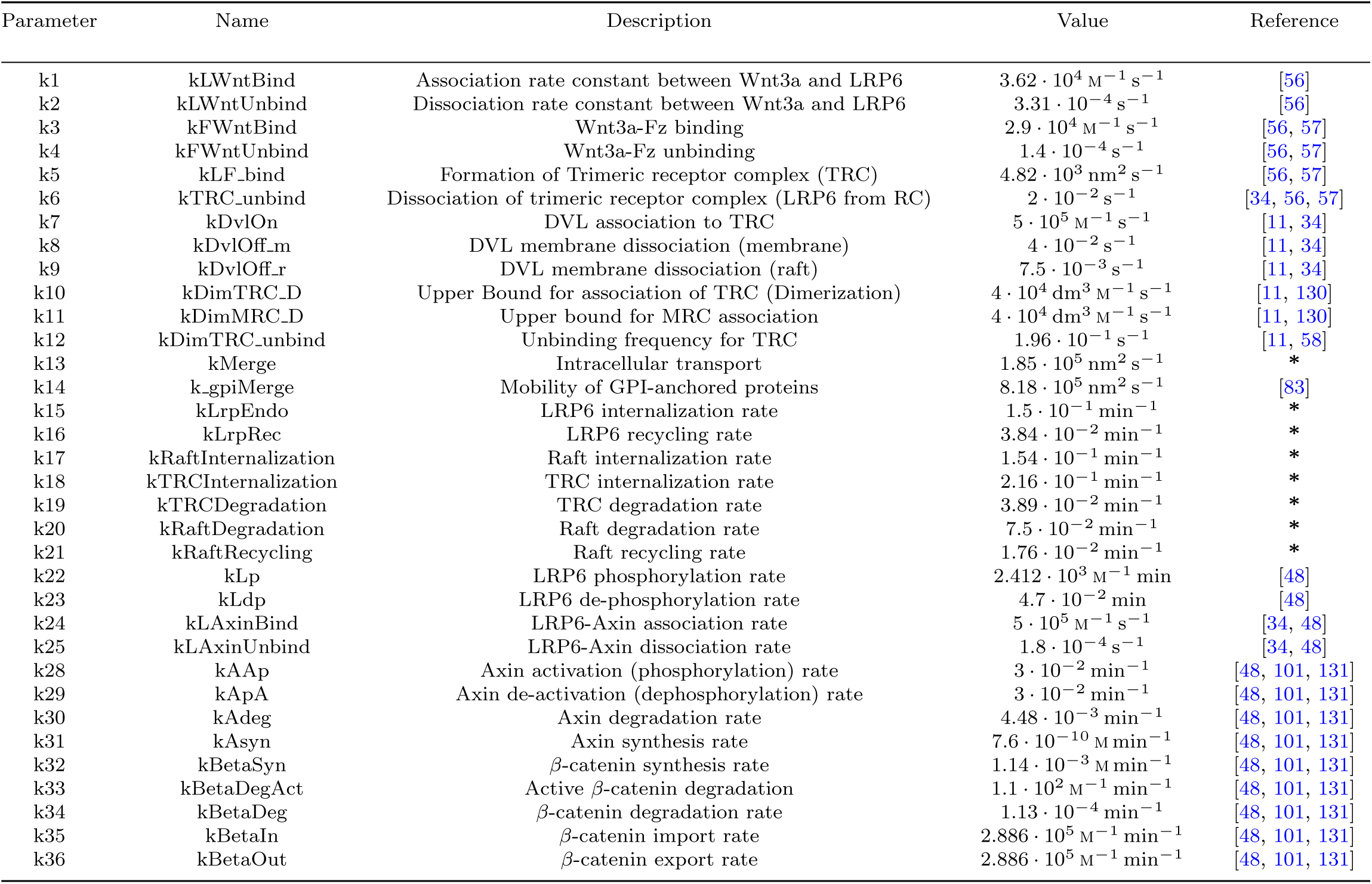
Parameter table containing all kinetic rate constants used in the model. Parameter names listed in column 1 correspond to the annotations in Figures 1 and 2. Parameter values that have been calibrated are marked with a star (*)

#### Wnt receptor binding (A)

The first step in the initiation of the signal transduction process is the binding of the ligand to its corresponding receptor. In the case of canonical Wnt signaling, signaling is initiated by the binding of specific Wnt proteins, such as Wnt3a or Wnt9, to membrane-associated LRP5/6 and Frizzled receptors. Numerous studies show that, under normal conditions, Wnt binding to both receptors, Frizzled and LRP6, is required to initiate canonical Wnt signaling [56–58]; and that binding to both receptors occurs simultaneously and independently [59]. Accordingly, the formation of the resulting trimeric ligand receptor complex (TRC) is modeled as a two-step process. First, extracellular Wnt binds to LRP6 or Fz. Subsequently, Wnt/FZ complexes bind to LRP6, or Wnt/LRP6 complexes bind to FZ. All processes are parameterized with the corresponding association and dissociation rate constants found in the literature [7, 56].

#### DVL binding (B) and receptor complex oligomerization (C)

Importantly, recent findings suggest that a single trimeric receptor complex (TRC) does not suffice for pathway activation. Instead, activation requires the oligomerization of at least two trimeric receptor complexes, mediated by the cytoplasmic protein Dishevelled (DVL) [11, 59, 60]. Consequently, we include the recruitment of DVL to the membrane and its binding to TRC and assume that the experimentally determined rate of Dvl self-aggregation serves as an upper limit for the dimerization/oligomerization rate of receptor complexes bound to DVL [10, 11, 58]. In this step, we take into account the varying membrane dwell times of DVL, which crucially depend on the localization of Frizzled [11] and the local membrane composition, in particular the PI(4,5)P2 density, which is typically elevated in lipid rafts domains [12].

#### Receptor phosphorylation (D) and signalosome formation (E)

DVL binding and receptor oligomerization prime the phosphorylation of multiple PPP(S/T)P motifs within the intracellular domain of membrane-anchored LRP6. In our model, we focus on the phosphorylation of site T1479 by membrane-associated CK1*γ*, which is pivotal for subsequent signal transduction of canonical Wnt signaling [13, 14, 61, 62]. Phosphorylation of site T1490 within the membrane-proximal motif by membrane-associated GSK3*β* is less specific and primarily serves to prime the CK1*γ*-mediated phosphorylation of site T1479 [13, 18]. Importantly, this process is limited to cholesterol-rich liquid ordered (lipid raft) domains [14, 15, 32, 63]. Since only a minor fraction of LRP6 receptors are located in lipid rafts [32, 34, 35, 64], the membrane localization of the receptor complex represents one of the most rate-limiting properties in LRP6 phosphorylation and the activation of the receptor complex.

#### Receptor life cycle

The dynamics of the receptor life cycle, including multi-complex formation, ligand-induced or raft-induced internalization, recycling, and intracellular sorting, represent important yet challenging features to incorporate into a computational pathway model. Together with the resulting heterogeneous distribution of signaling components, such as receptors and kinases, these dynamics play a decisive role in canonical Wnt signaling. Here, we use a compartmental approach to represent cellular components (membrane or nucleus) and subcellular or membrane-based components (raft domains, endosomes, lysosomes) as individual entities. These entities may contain variable amounts of molecules (or other compartments) and may have arbitrary attribute values to represent important characteristics of the compartment, like its radius or volume. The model formalism used here, ML-Rules, enables the dynamic creation, modification, and removal of compartments during runtime [65, 66]. Processes like internalization, recycling, merging, or lysosomal degradation can be expressed straight-forwardly in terms of reaction rules. Corresponding attributes, such as radius or volume, can be dynamically combined, modified, or newly specified. More importantly, these dynamic attributes are available for reaction rate calculations, allowing the specification of individual volume-dependent reaction rate kinetics [53]. Species, such as molecules, located within a compartment, are handled according to the given rule. This means, for example, in the case of internalization, they are transferred from one compartment to the other, and in the case of compartmental degradation, all molecules within the compartment are removed from the system. Based on this, we were able to incorporate the essential processes of the LRP6 life cycle during canonical Wnt signaling in our computational model, as depicted in Figure 2.

**Fig. 2:**
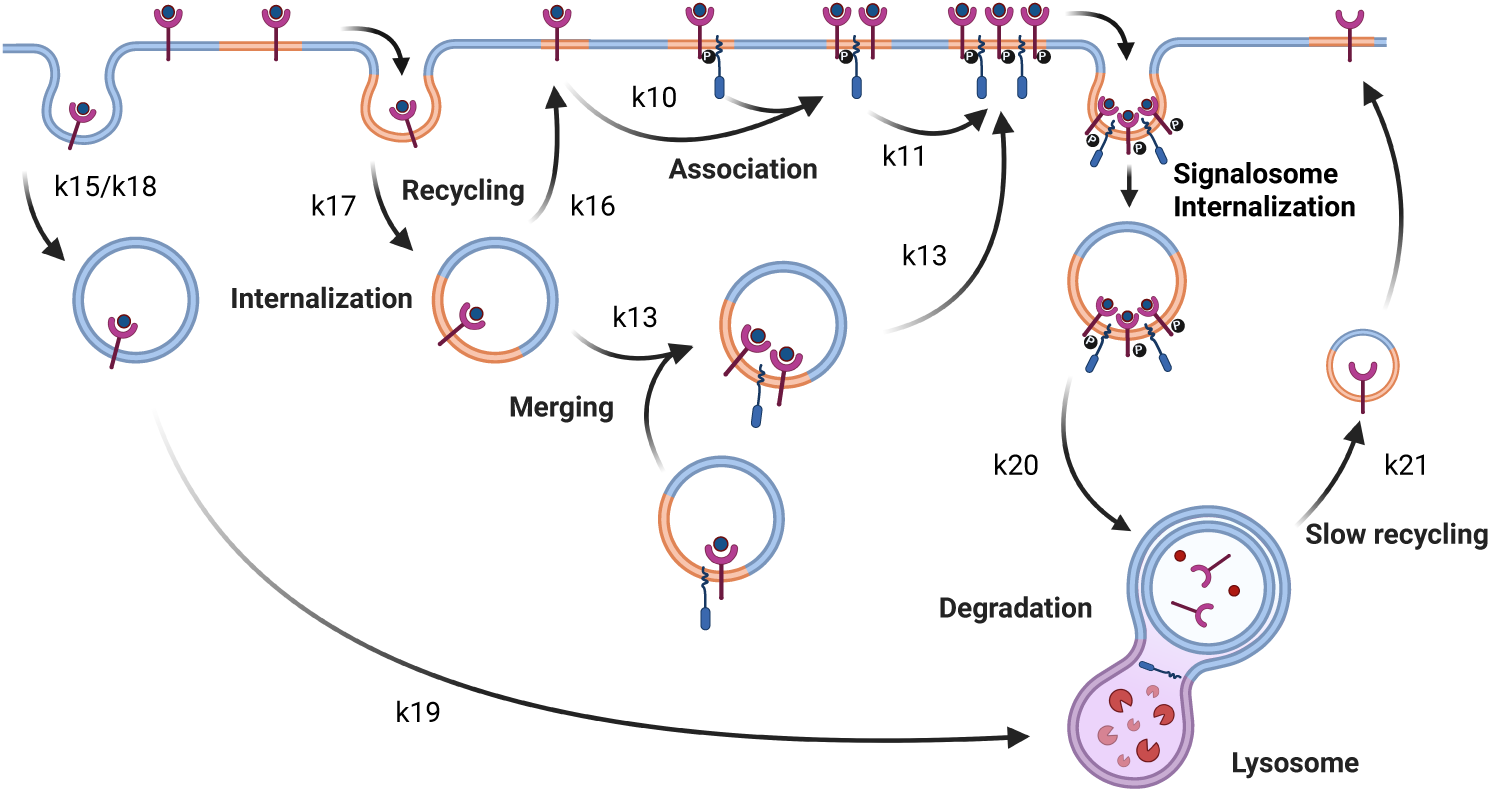
Simplified view of receptor life cycle as implemented in our model. For better accessibility, only LRP6 is considered; its binding state to FZ and cytosolic proteins, such as Dvl or Axin, are omitted in the figure. Variable names of kinetic rate constants are associated to each arrow representing rules implemented in the model. Created in BioRender. Haack, F. (2025) https://BioRender.com/nyn65rn

#### Downstream Signaling

Once phosphorylated, the intracellular domain of LPR6 within the multimeric receptor complex (MRC) provides high-affinity binding sites for Axin. This induces the recruitment and binding of Axin and other components of the *β*-catenin destruction complex to the receptor complex at the membrane. This leads to the inhibition of the destruction complex and allows for *β*-catenin aggregation in the cytosol and nucleus. Reactions involving the interaction of LRP6 and Axin and subsequent downstream signaling events are part of a model that has previously been published and is being reused here [48].

### 2.2 Data-driven model building unravels three concurrent mechanisms driving receptor internalization in the initial phase of Wnt signaling

Important steps in the initial phase of canonical Wnt signaling are the formation of the trimeric Wnt/Fz/LRP6 receptor complex, i.e. the binding of the extracellular ligand Wnt to its cell surface receptors LRP6 and Frizzled, their subsequent association. The binding kinetics between Wnt and its pathway-specific receptors LRP5/6 and Frizzled receptor subtypes, as well as the molecular structure of the Wnt/Fz/LRP6 complex, have been well characterized experimentally [7, 56, 67]. However, the localization of the Wnt-receptor interaction, the impact of the membrane environment, and the role of posttranslational lipid modification of Wnt are still under discussion [68, 69]. Our quantitative, mechanistic modeling approach allows us to explore these processes in detail and provide quantitative evaluations of common theories, e.g., why Wnt is primarily located in ordered membrane domains, and what role lipid modifications of Wnt proteins, such as palmitoylation, play in this context.

While direct experimental observations under endogenous conditions are barely available, time-resolved experimental measurements on receptor dimerization [70, 71] and internalization (e.g. [32, 71–73]) provide valuable kinetic data. This data is used to evaluate different model scenarios with respect to the localization of Wnt and its binding to receptors FZ and LRP6. First, we consider LRP6 endocytosis induced by MESD, a specific ligand that binds LRP6 and induces its internalization. As MESD binds to LRP6 independently of its localization at the cell surface, kinetic rates for the internalization and recycling of LRP6 in response to unspecific MESD-LRP6 binding provide an upper bound for the internalization of LRP6 [72, 73]. The internalization of LRP6 in response to MESD treatment can be reproduced using a simple model comprising only three rules: MESD/LRP6 binding, LRP6 internalization, and LRP6 recycling.

In contrast to MESD, several studies report that Wnt binding to LRP6 selectively occurs in ordered membrane environments, where it gets internalized with the resulting receptor complex [15, 68, 69]. We performed several simulation experiments with different assumptions regarding the spatial restrictions of the Wnt/LRP6 interaction and the order of events leading to the formation of the trimeric receptor complex, consisting of Wnt/FZ/LRP6. For each assumption, we tried to fit the corresponding model to available experimental data on receptor dimerization and internalization. Intriguingly, our simulation results (Figure 3) indicate that Wnt binding to LRP6 in ordered domains alone does not quantitatively suffice to reproduce the internalization and receptor complex dimerization rates observed in vitro (e.g. [32, 71]). This can be explained as follows: with LRP6 being homogeneously distributed throughout the membrane, only a minor fraction (25-30 %) is located in ordered membrane environments [15, 32, 63, 68] and this distribution does not significantly increase after Wnt stimulation [32, 63, 71]. Therefore, only a small fraction of LRP6 is available for Wnt binding and subsequent internalization, which is in contrast to most experimental measurements, indicating the internalization of large fractions of cell surface LRP6 (50-60%) within several minutes after Wnt stimulation [32, 73, 74]. Thus, according to our results (Figure 3), LRP6 internalization is orchestrated by three different mechanisms: Wnt/FZ/LRP6 complex internalization outside membrane domains, direct LRP6 internalization, and signalosome internalization in ordered membrane domains. Together, these concurrent processes yield the correct amount/fraction of internalized LRP6.

**Fig. 3:**
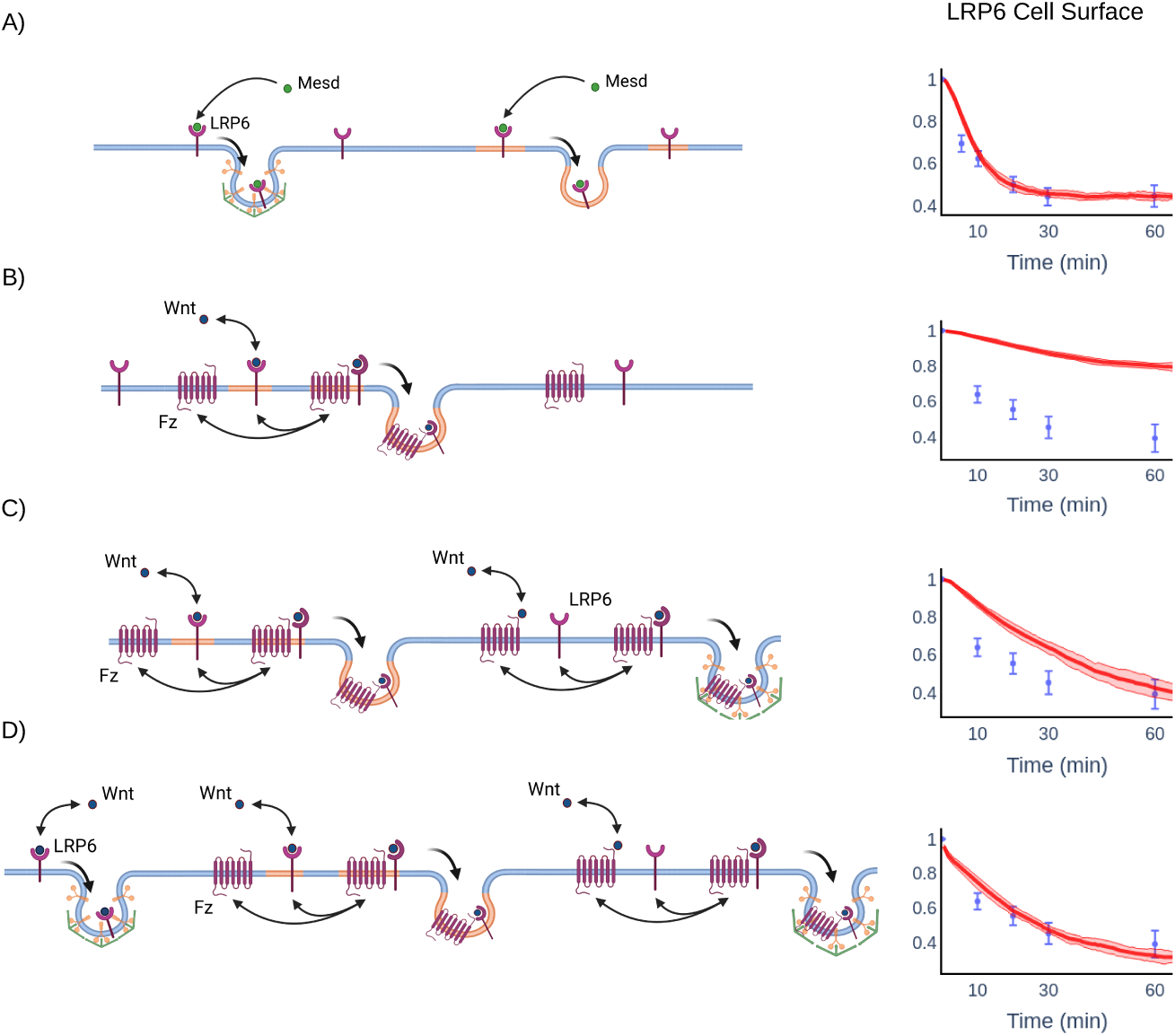
Comparing LRP6 internalization for different model scenarios. A) Simplified model representing MESD-induced internalization of LRP6. The trajectories exhibit a good fit to the experimental values measured in [73]. B) More complex model, in which Wnt binding is restricted to LRP6 in lipid raft domains. No binding occurs between Wnt and Frizzled. Apparently, the simulation result of the model with the best fit cannot reproduce the experimental data as measured, e.g. [32]. C) Complex model, where Wnt binding to LRP6 is still restricted to raft domains, but FZ-Wnt interaction is allowed everywhere. D) Complex model, where Wnt binding to LRP6 and Frizzled is allowed everywhere. Created in BioRender. Haack, F. (2025) https://BioRender.com/7htzq6h

### 2.3 Microdomains reinforce rebinding of DVL, which is crucial for the formation of multimeric receptor complexes

The localization of the Wnt/FZ/LRP6 complex (TRC) to ordered raft domains is extremely important for the stability of the receptor complexes, as shown in the next simulation experiments. Several studies indicate that the membrane affinity of DVL, and hence its membrane dwell time, highly depends on the membrane composition [12, 34, 35].

A model that incorporates the concept of rebinding (the process by which proteins return to their original location) and residence time (the duration a protein remains in a particular location) can explain how compartmentalization, induced through lipid rafts and membrane cytoskeleton facilitate the DVL-mediated dimerization of the TRC, which is crucial for Wnt signaling.

To illustrate this, we consider a small part of the entire model and use the resulting submodel for a set of simulation experiments. The simplified model contains a fixed number of established trimeric receptor complexes (TRC), which are either located in the membrane or in membrane sub-compartments representing ordered membrane domains, and an intracellular pool of DVL molecules that can associate with TRC. The residence time of DVL at the membrane, i.e., its dissociation rate from the receptor complex, depends on the localization of the TRC, as observed in several in-vitro studies [11, 12, 34]. As a result of the lower dissociation rate of the DVL-TRC complex in raft compartments, the equilibrium number of DVL bound to the receptor complexes in rafts is similar to the number of DVL-TRC complexes in the remaining membrane compartment – despite the fact that only 25% of TRCs are located in raft compartments (red line vs brown line, preliminary Fig. 4). This effect is even stronger for the dimerization of the receptor complexes, mediated by DVL self-aggregation (purple vs blue line, preliminary Fig. 4), as only in raft compartments a stable population of DVL-TRC dimers is established. This is due to the low self-aggregation rate of DVL, which controls the dimerization of the TRC [10, 62, 75].

**Fig. 4:**
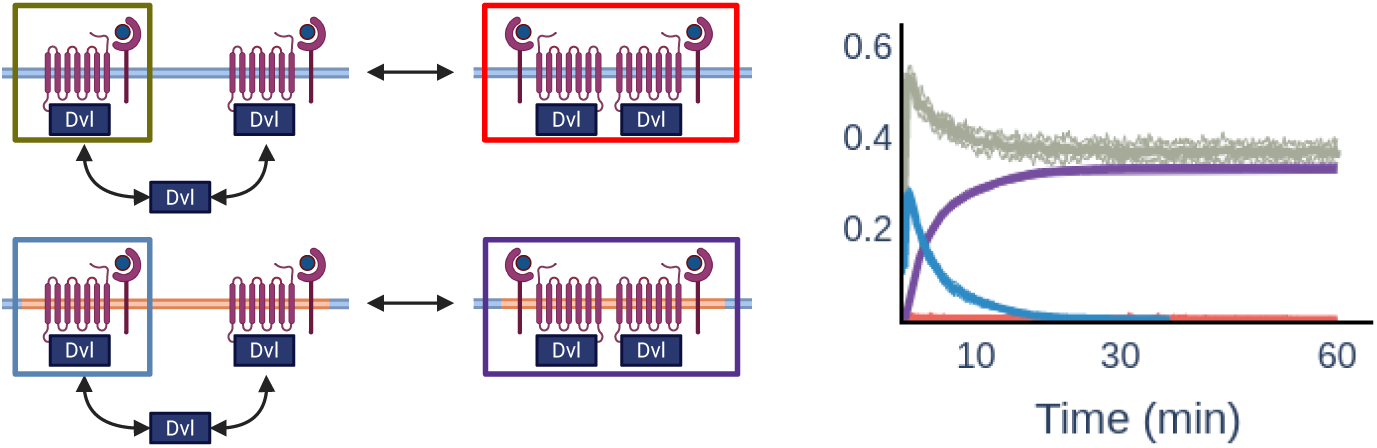
Simulation of sub-models containing a fixed number of trimeric receptor complexes (TRC), to which cytosolic DVL may bind or unbind. TRC with bound DVL form dimers. Colored boxes in the left figure correspond to the observed model entities, whose simulation trajectories are displayed on the right-hand side. Simulation trajectories indicate almost exclusive accumulation of DVL/TRC dimers in raft domains (purple) compared to non-raft regions (red). This due to the small volume of raft compartments and longer membrane residence time of DVL, facilitating immediate binding and rebinding of TRC located in the same raft compartment. Created in BioRender. Haack, F. (2025) https://BioRender.com/f5qyuaz

Due to the small volume of each individual compartment, the chance of immediate rebinding between DVL/TRC complexes before DVL disengages from the membrane is much higher than in the remaining (large) membrane compartment. This leads to a high (equilibrium) number of dimerized TRC in ordered membrane domains and a very low number of dimerized TRC in the remaining membrane. Thus, the simplified model provides an illustrative explanation of why aggregates of DVL and TRC are primarily located in ordered membrane domains and highlights the crucial role of these raft domains in recruiting and binding DVL to the receptor complex, leading to the formation of the signalosome.

### 2.4 Cross-compartmental mobility of membrane proteins is essentially required for the formation of large signalosomes due to sparse receptor density

After binding of pathway-specific Wnt proteins, the canonical Wnt signaling pathway requires the co-localization of LRP6 and Frizzled, subsequent receptor clustering, and signalosome formation to activate signal transduction. For this, the lateral mobility of membrane proteins is essentially required. Proteins and lipids undergo 2-dimensional lateral diffusion within fluid biological membranes. High-speed single-particle tracking revealed that membrane proteins diffuse freely only within ∼40–250 nm voxel bounded by actin “fences” [76]. For longer distances (and observation time), membrane-bound proteins yield a characteristic hop-diffusion behavior with effective diffusion coefficients being significantly lower compared to lateral diffusion rates within individual voxels [77, 78]. It is widely accepted that actin-associated scaffolds (for example, ERM proteins linking lipid rafts) can pre-cluster receptors by transiently co-anchoring them, and that the confinement through the cytoskeleton in general increases local receptor density and rebinding of ligand or effector molecules accelerating oligomerization (e.g., EGFR dimerization) and downstream signaling [77, 79]. This presupposes that several pathway-specific membrane-anchored proteins or protein complexes are located in the same or in neighboring voxels. However, a quantitative consideration of the typical density of cell surface receptors compared to the size of voxels bound by actin “fences” reveals a surprisingly large separation of receptors: Assuming a mean cell surface of a medium-sized cell, such as HeLa cells (1, 5*x*10^3^*µ*m^2^)[80] and the median size (length) of the cytoskeletal compartment in HeLa cells of 64nm [78, 81], then an individual voxel covers 4096nm^2^ or 4, 096 × 10*^−^*^3^*µm*^2^. Projected to the entire cell surface, more than 360000 voxels span the cell surface. In contrast, typical numbers of cell surface receptors range between 1k and 10k. This means that, for homogeneously distributed receptors, there are not several receptors within one compartment; instead, single receptors must surpass tens or hundreds of actin “fences” to co-localize in the same compartment and interact. This issue arises as soon as the receptors form heteromeric receptor complexes. The density becomes even sparser, while confinement within a compartment increases with complex size, eventually leading to temporary immobilization within a membrane compartment. This is exactly the case in canonical Wnt signaling. Canonical Wnt signaling depends on the formation of a large receptor-protein complex, termed signalosome, initiated by co-localization of its pathway-specific receptors LRP6 and FZ after Wnt binding and subsequent recruitment of cytosolic proteins, such as DVL, Axin, and casein kinases. For successful receptor activation upon Wnt binding, Frizzled, LRP6, and Wnt need to form a trimeric receptor complex (TRC), followed by the recruitment and binding of cytosolic DVL (D-TRC). Recent studies indicate that at least two of these receptor protein complexes are needed for a multimeric receptor complex (MRC) [59, 67] and that receptor clustering is required for successful pathway activation [82]. Considering the complex mechanisms and large receptor protein complexes required to activate signal transduction in canonical Wnt signaling [82], we next examine whether signalosome formation is achievable just by passive lateral diffusion or whether active mechanisms are additionally required to support receptor aggregation and signalosome formation. Experimental measurements show that receptor cluster size in canonical Wnt signaling can reach up to ∼100 nm and more [83]. As outlined before, larger protein complexes are located in distinct membrane compartments surrounded by an actin skeleton and protein “picket” fences, hence mostly immobilized when crossing a certain size threshold (*>*20 nm) [83–85]. In canonical Wnt signaling, this upper threshold roughly applies to dimerized TRC and any larger receptor complex, hence all species we consider as multi receptor complex (MRC), are above that limit, hence are expected to be largely restricted in their lateral mobility.

Therefore we analyze the impact of immobilization of multimeric receptor complexes (MRC) on the formation of signalosomes. We consider model scenarios in which single TRC are always mobile, while higher-order receptor complexes (MRC) are mobile or immobile for varying Wnt stimuli. In the case of immobile MRC, larger receptor complexes emerge only by the lateral association of TRC with existing MRC. Fig. (5) shows the result of our simulation as the number of TRC molecules located in signalosomes for low, medium, or high Wnt stimuli at 3 hours after the onset of WNT stimulation. In the model scenario, where MRC are mostly static and form immobile clusters, as suggested by the literature (e.g. [83, 86]), we find that only a small amount of medium-sized and no large receptor clusters emerge, even for high Wnt stimuli (see Fig. 5).

**Fig. 5:**
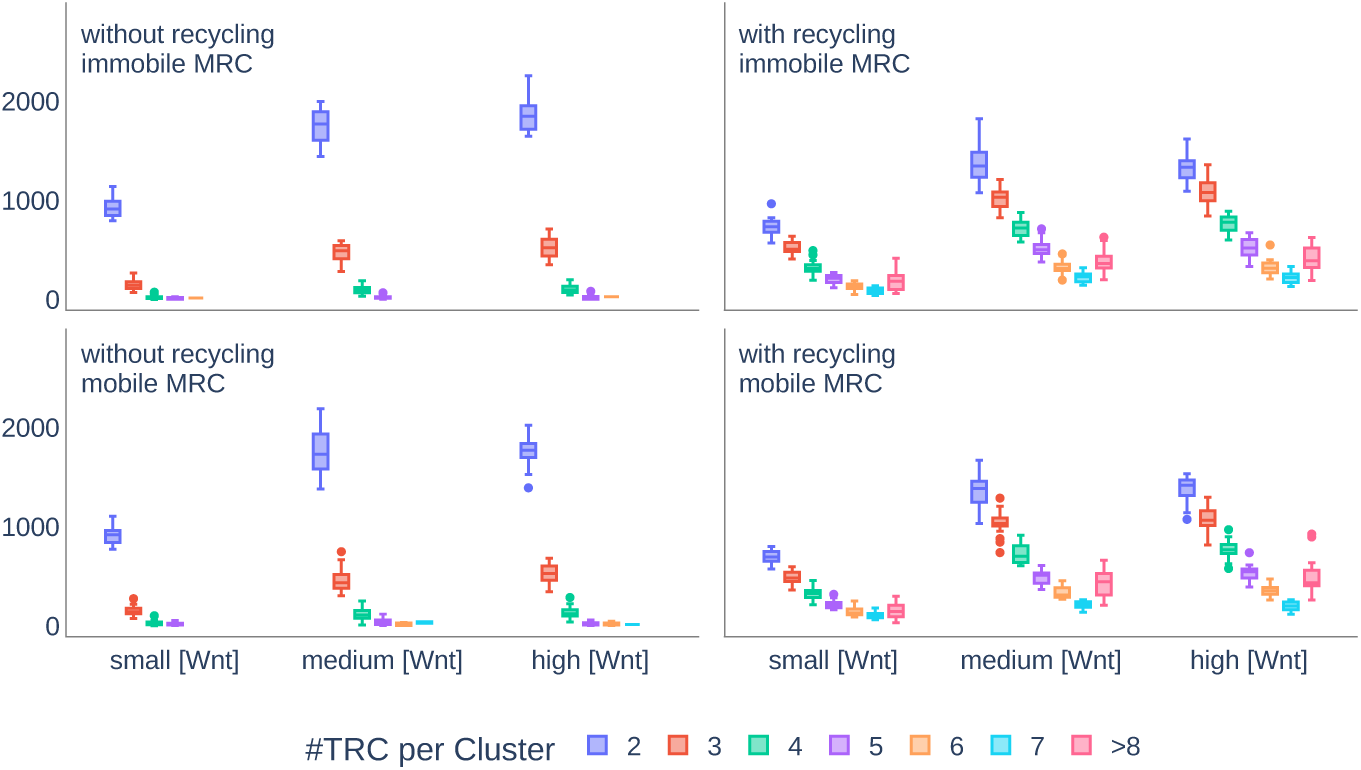
Size distribution of receptor complexes in varying model scenarios. Considered are model variants with and without fast receptor recycling (columns) and model variant with or with receptor cluster immobilization (rows). Size of the receptor clusters (signalosomes) is estimated by the number of Wnt/Fz/LRP6 receptor complex (trimeric receptor complexes – TRC) inside an individual signalosome. Number of signalosomes (Y-axis) is calculated by the sum over all timepoints between 1h and 6h of an individual simulation trajectory and averaged over all replications. In a model including internalization and fast recycling the amount of medium and large receptor complexes is largely increased, indicating that these processes are required for the formation of large signalosomes.

Surprisingly, a model scenario including mobile MRC does not lead to a higher amount of large signalosomes either. We thus hypothesize that an additional mechanism is required for the formation of large signalosomes, which we will explore in the following.

### 2.5 Receptor trafficking, endosomal sorting, and fast recycling promote formation of large signalosomes

Numerous studies have suggested a beneficial role of receptor trafficking in various signaling pathways [36, 87, 88] However, despite the important role of internalization in Wnt signaling, and our simulation experiments have confirmed this, few studies exist that have actually analyzed the potential role of receptor trafficking and recycling in Wnt signaling [16, 89]. Therefore we ask whether receptor trafficking, endosomal sorting, and recycling might also play a beneficial role in Wnt signaling, as recently discussed for other signaling pathways.

In our model we differentiate between raft-associated and non-raft-associated internalization. Endocytosis of receptor or receptor complexes bound to Wnt outside of raft domains is most likely mediated by ZNRF3 and RNF43, leading to degradation of the receptor complex and slow recycling [20]; whereas endocytosis of raft-associated receptor or receptor complexes bound to Wnt is subject to intracellular transport, hence quickly recycled back to the membrane, where they merge with existing raft domains [90–92]. Receptor complexes that are phosphorylated and in complex with members of the destruction complex (in our model simply represented by AXIN species), hence fully assembled signalosomes, are transferred to the lysosomal degradation pathway upon internalization. The associated receptors LRP6 and Fz are recycled back to the membrane as single species (see Fig. 2).

Intriguingly, activating receptor trafficking mechanisms in our model yields numerous medium-sized and even large signalosomes, as observed in vitro [14, 82, 83, 93] (Fig. 5). This result can be explained as follows: According to in-vitro data and the results of our in-silico study, Wnt/Fz/LRP6 receptor complexes, either monomeric, dimerized, or as part of a larger complex, are rapidly internalized after WNT binding. At the same time, the aggregation rate between these complexes is comparatively low [11]. Therefore TRCs are typically internalized before they can co-localize and aggregate with other TRC or existing MRCs, which limits the aggregation of TRC into higher order receptor complexes. This effect is counteracted by receptor trafficking and immediate/fast recycling of TRCs.

Further we can assume that the mobility of receptor-bearing endosomes is higher than lateral mobility at the membrane. Note that this aspect becomes even more relevant considering our quantitative analysis of receptor density and compartmentalization of the membrane (see above). While lateral diffusion of the receptor complexes is likely subject to local immobilization and hop diffusion, receptor-bearing endosomes might further be driven by active, intracellular transport, which potentially increases the mobility of receptor complexes located in early endosomes. This effect has not been considered in our model, but might increase the observed promoting impact of receptor trafficking on signal transduction. In addition, active sorting has been observed for endosomes with high content of saturated lipids and cholesterol; studies show a gradient of lipid composition, which shifts from saturated to unsaturated lipids with increasing distance from the cell surface [94]. This means, in contrast to pure lateral association of TRCs and MRCs, the internalization and recycling support and enhance the localization of receptor complex components, such as LRP6, Frizzled, and Dvl, in microdomains that are enriched in cholesterol and saturated lipids [92]. In summary, to arrive at the size of large receptor clusters as described in the literature, an active sorting mechanism is required, most likely mediated, or at least supported, through endocytosis and fast recycling.

### 2.6 Lack of fast recycling attenuates LRP6 dimerization and phosphorylation leading to significantly reduced *β*-catenin accumulation

Our simulation results reveal a promoting effect of receptor trafficking on receptor clustering and signalosome formation. In the literature, increased receptor clustering and larger signalosomes typically imply an improved receptor activation as well as higher downstream signal output. To check whether this also applies to our model, we analyze the impact of receptor trafficking on LRP6 phosphorylation, and eventually on signal transduction in terms of *β*-catenin aggregation. Figure 6 shows the mean trajectories of model simulations with and without receptor trafficking in response to a medium Wnt stimulus.

**Fig. 6:**
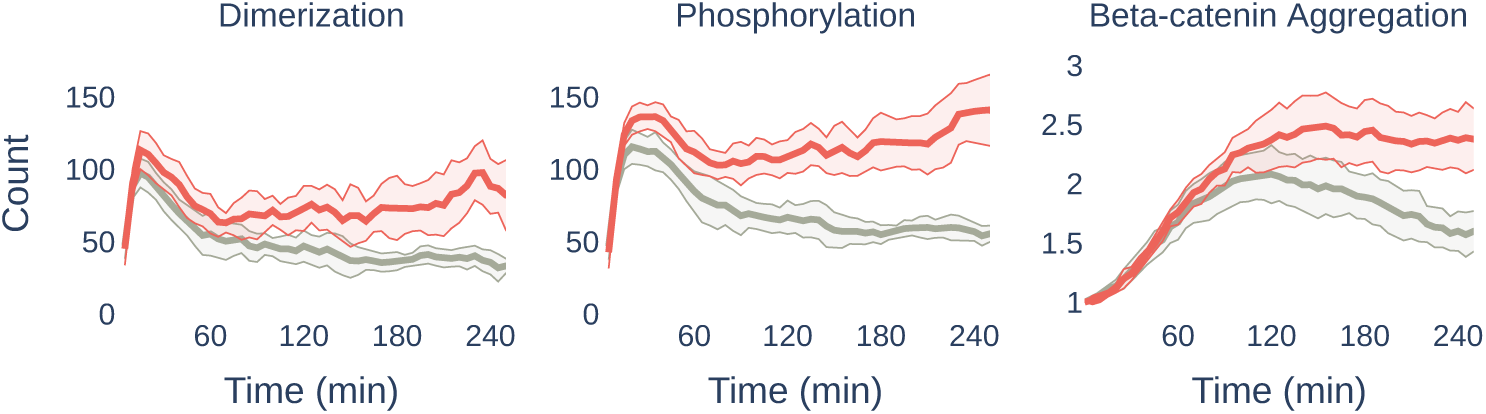
Mean simulation trajectories for receptor dimerization (left), phosphorylation (middle) and nuclear *β*-catenin concentration (right, fold change). The red line corresponds to simulation trajectories of a model with receptor trafficking and fast recycling. The grey line shows simulation trajectories of a model missing fast receptor recycling. Simulation results suggest, that the lack of fast recycling attenuates TRC dimerization and phosphorylation, leading to reduced *β*-catenin signaling.

As expected, the higher amount of larger signalosomes obtained in the model with receptor trafficking also yields a higher amount of phosphorylated LRP6 receptors as well as *β*-catenin counts in the nucleus. This indicates an attenuated pathway activity in a model lacking intracellular receptor trafficking. Interestingly, the deviation between the two scenarios (receptor trafficking vs. pure lateral diffusion) is small in the very early response after the onset of Wnt stimulation, but strongly increases after several minutes. The reason for this is that dimerization of TRC is sufficient for LRP6 phosphorylation and downstream signaling, a process achieved and primarily controlled by lateral diffusion directly after the onset of Wnt stimulation [11, 95]. However, TRC and dimerized receptor complexes are both subject to internalization. In the model scenario without fast recycling (gray color in Figure 6) all receptor complex entities are primed for degradation and slow recycling. In contrast, in a model with fast recycling raft-associated receptor complexes (mono and dimers of TRC) are recycled back to the membrane within several minutes, yielding larger and more stable signalosomes, that amplify LRP6 phosphorylation, and *β*-catenin stabilization, hence downstream signaling. This is in line with the general time frame typically obtained for fast recycling processes, which are in the range of ∼ 3-5 minutes [41, 96, 97].

### 2.7 Sustained, long-term Wnt signaling depends on internalization and receptor recycling

Notably, comparing model scenarios with and without receptor trafficking revealed that the effect of receptor trafficking not only transiently promotes the signal transduction, but also leads to a more sustained, long-term stabilization of *β*-catenin levels. Sustained, long-term activation of the Wnt/*β*-catenin pathway is mandatory for numerous physiological processes, such as embryonal patterning and morphogenesis (e.g. [98]), adult stem-cell maintenance and self-renewal (especially in intestinal crypts) (e.g. [1, 4]), and tissue regeneration (e.g. [3]). At the same time sustained, long-term Wnt/*β*-catenin is an important hallmark of Wnt-induced cancer [99].

To analyze the impact of receptor trafficking on long-term Wnt signaling and for a more detailed model exploration, we employ a recent visual analytics approach [100]. In this visualization design, time series plots are embedded in parallel coordinates, enabling a simultaneous and interactive visual analysis of numerical model parameters and simulated signaling behaviors over time. In the visualization, each line in the graph corresponds to an individual model configuration that has been simulated in the course of a parameter scan (see Figures (7 and 8). The gray parallel axes depict the individual parameter values employed in the parameter scan (e.g., kRaftInternalization, kRaftRecycling, kRaftDegradation), whereas the colored axes depict simulation trajectories of a certain model observable, such as LRP6 internalization (LRP6Int) or nuclear *β*-catenin levels clustered into groups of similar time series.

The visualization allows for exploring the relation between certain model configurations and model outputs in more detail. For instance, Figure 7 shows two noteworthy model configurations, highlighted in orange and blue, where the simulation trajectories show very different model outcomes in terms of LRP6 dimerization (blue axis), phosphorylation (red/orange axis), and *β*-catenin accumulation (violet axis), despite similar parametrization and Wnt stimulus (nWnt). The only parameter values that differ significantly between the model configurations are related to fast and long-term recycling (kMerge and kRaftRecycling), which supports our previous observation that receptor trafficking has an unexpectedly strong impact on *β*-catenin aggregation.

**Fig. 7:**
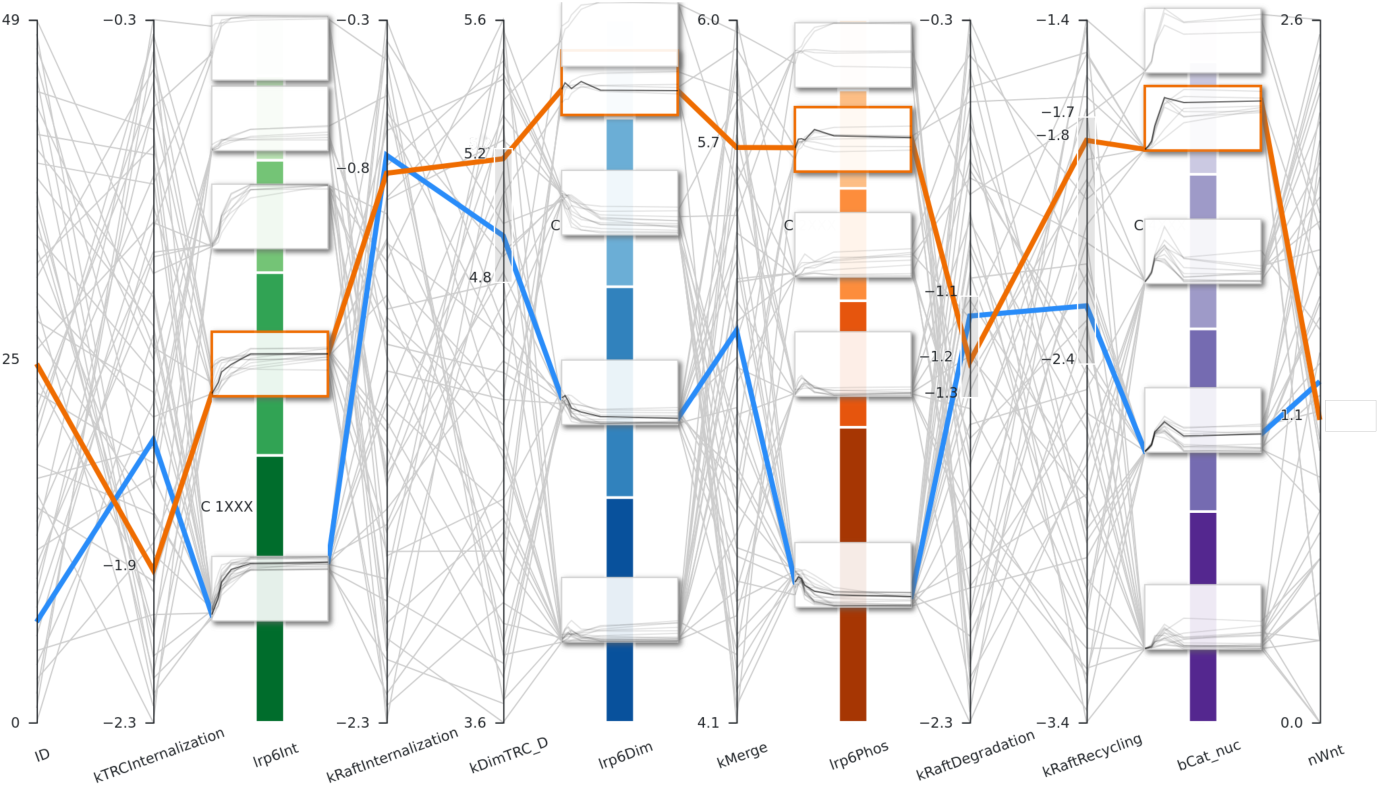
Parallel coordinates depicting parameter values of various model configurations (lines) and corresponding simulation results (boxes of clustered trajectories). Two model configurations highlighted in the figure by orange and blue lines illustrate the impact of varying the parameter values for fast and slow recycling (kMerge, kRaftRecycling) on output variables, such as phosphorylation in red, LRP6dimerization blue, and in particular on *β*-catenin aggregation (shown in purple).

Next, we ask which parameter value combinations are required for the optimal – i.e., strongest – activation of the pathway. For this, we consider all simulation trajectories with the highest and most stable *β*-catenin accumulation. These trajectories are displayed in the uppermost group of the *β*-catenin axis in Figure 8. By selecting this group we highlight all model configurations that yield the corresponding simulation trajectories (blue lines).Intriguingly, model configurations that result in the most efficient signaling outcome have a clear pattern and are characterized by high values of slow receptor recycling (kRaftRecycling) and moderately high values of parameters associated with receptor internalization and fast recycling (kRaftInternalization, kMerge). Extending the selection to the second group of high, stable *β*-catenin (black lines, Fig. 8) results in a more diverse parameter value distribution, which means that not every parameter value of receptor recycling has to be high. The only exception is the parameter value of slow receptor recycling (kRaftRecycling) and to some extent fast receptor recycling (kMerge), which are essentially required to be high for long-term *β*-catenin stabilization.

**Fig. 8:**
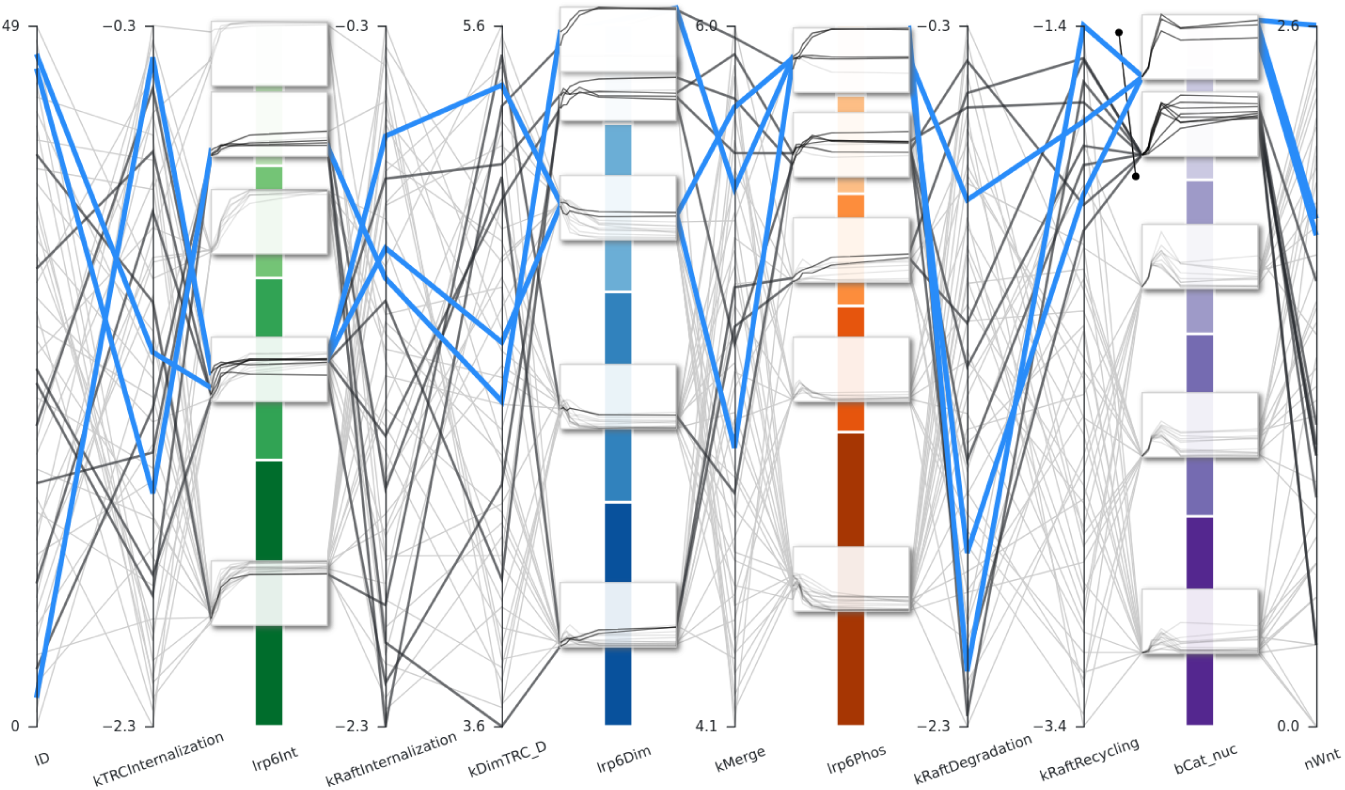
Parallel coordinates depicting parameter values of various model configurations (lines) and corresponding simulation results (boxes of clustered trajectories). In the figure all model configurations are highlighted, that lead to high and stable *β*-catenin aggregation, for high and medium Wnt stimulus.

**Fig. 9:**
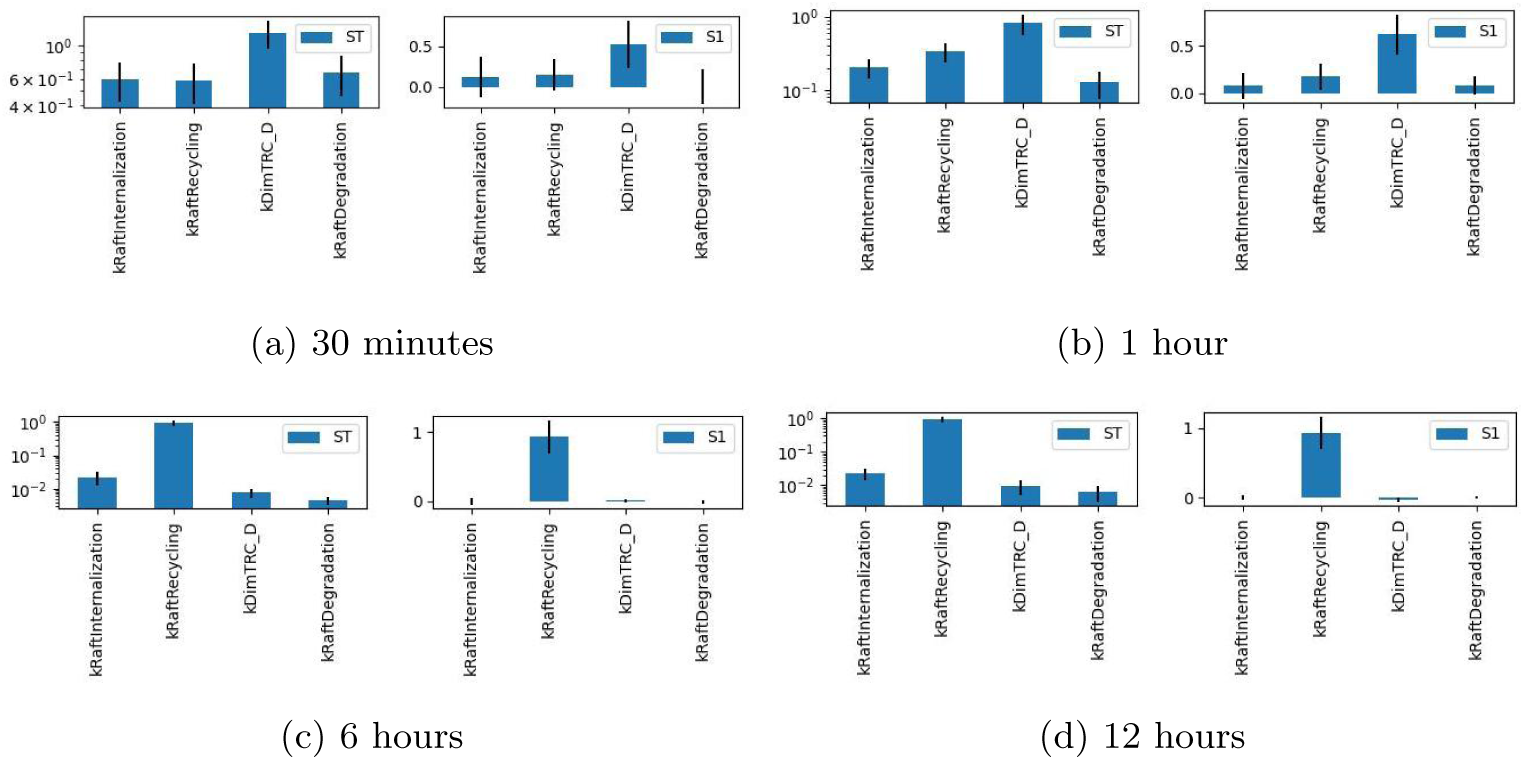
Global (ST) and first-order (S1) Sobol indices measuring model sensitivity (in terms of *β*-catenin concentration) against changes in receptor trafficking dynamics measured at different time points after onset of WNT stimulation. Parameters with high Sobol indices differ between early, transient (30 minutes and 1 h) and late, long-term response (6h and 12h) indicating that *β*-catenin stability is controlled by different mechanisms in a time-dependent manner.

To challenge our previous insights drawn from the visualization and to quantify how changes in the regulation of membrane dynamics alter the signal outcome hours after signal initiation, we apply a global sensitivity analysis using the Sobol’ method. We specifically vary the parameter values of all processes related to receptor trafficking and observe *β*-catenin accumulation in the nucleus at different time points after signal initiation (30 minutes, 1 hour, 6 hours, and 12 hours). As a result of the sensitivity analysis, we observe a surprisingly strong difference in Sobol indices between early, transient (30 minutes and 1 h) and late, long-term response (6h and 12h). For early time points, the kinetic rate for trimeric receptor complex dimerization driven by lateral diffusion (kDimTRC D) has the highest Sobol indices (global (ST) as well as first-order (S1)), followed by parameters related to receptor trafficking (kRaftInternalization, kRaftRecycling, kRaftDegradation). This is in line with our previous results. For later time points, however, the recycling rate (kRaftRecycling) of internalized and degraded receptor complexes is the dominating parameter that controls *β*-catenin levels. These results imply that different mechanisms control *β*-catenin stability, hence pathway activity, in a time-dependent manner and that recycling plays a surprisingly vital role in long-term *β*-catenin signaling.

To explain these results, we need to consider the signaling logic of canonical Wnt signaling, which is fundamentally distinct from other prominent signaling pathways, such as EGFR or TGF-*β*. Instead of inducing the intracellular signal transduction by G-proteins or kinase cascades, canonical Wnt signaling is regulated post-translationally by stabilizing the pathway’s target protein *β*-catenin. This is achieved by the recruitment and binding of essential components of the destruction complex, which limits *β*-catenin levels by continuous degradation, to the multimeric receptor complex (MRC) that forms upon binding of Wnt to its co-receptors Frizzled and LRP5/6 and subsequent receptor clustering (as described in previous sections). Since the signalosome lacks downstream enzymatic amplification steps, signal persistence depends solely on ongoing ligand-receptor clustering and the inhibition of *β*-catenin degradation, rather than second-messenger kinetics.

Here, the scaffold proteins AXIN/AXIN2 play a crucial role. AXIN is the rate-limiting scaffold of the *β*-catenin destruction complex because it is present at very low cellular concentrations and serves as the central platform that recruits all other key components (APC, GSK3, CK1) [101, 102]. Upon Wnt activation, AXIN is rapidly recruited to LRP5/6 and is co-internalized with the receptor complex into caveolin-enriched endocytic vesicles / multivesicular bodies (MVBs), where destruction-complex components (AXIN, GSK3) become sequestered. Since AXIN availability dictates destruction complex formation, its depletion quickly induces *β*-catenin accumulation [101, 103]. However, the axin-related protein AXIN2 (also called Conductin) is a well-established transcriptional target of *β*-catenin/TCF complexes and is induced by active Wnt signaling [4, 17, 104]. Both AXIN proteins are functionally equivalent in vivo [105]. In contrast to AXIN1, which is constitutively expressed, AXIN2 expression increases following pathway activation and acts as part of a delayed negative feedback loop in canonical Wnt signaling [106–108].

Accordingly, the degradation of endogenous AXIN, induced by LRP5/6, has been considered to be a critical event in canonical Wnt signaling from early on [109–111] However, in view of our findings revealed by quantitative modeling, one important aspect has been neglected: If internalization and subsequent lysosomal degradation of the entire signalosome are the driving mechanisms for AXIN degradation, the recycling of LRP6 is essential to achieve a continuous receptor engagement of AXIN. Without a continuous round of receptor recycling, long-term Wnt signaling either leads to the depletion of cell surface LRP6 receptors, which lowers receptor density and signal activation efficiency (see above), or eventually all LRP6 receptors at the cell surface will be engaged in signalosomes, limiting the availability for newly synthesized AXIN2 proteins.

From a quantitative perspective, none of the known feedback mechanisms is capable of compensating for the negative feedback loop of AXIN/AXIN2, which is required for sustained, long-term Wnt signaling. While AXIN2 expression increases after pathway activation, the resulting protein does not uniformly restore *β*-catenin degradation because its stability and function are themselves regulated by Wnt-dependent post-translational mechanisms, including tankyrase-mediated turnover [112]. However, tankyrase expression is generally not transcriptionally co-regulated with *β*-catenin–driven AXIN2 expression, and neither shows a reliable correlation across tissues or experimental conditions. Moreover, AXIN2 participates not only in the cytoplasmic destruction complex but also contributes to the assembly of the membrane-proximal signalosome, thereby conferring context-dependent roles in both attenuating and facilitating Wnt pathway output [108, 113, 114].

Lastly, there is no experimental evidence that LRP6 expression is correlated with canonical Wnt activation; on the contrary, several studies report a negative-feedback regulating *β*-catenin through LRP6 desensitization and down-regulation of LRP6 expression [73, 74].

## 3 Discussion

In our simulation model, we combined a multitude of experimental observations into a coherent explanatory framework that also considers the quantitative properties and dynamics of the spatio-temporal regulation of canonical Wnt signaling. For this purpose, a new, optimized multi-level rule-based modeling and simulation approach is employed to describe and simulate the compartmental distribution of extracellular, intracellular, and membrane-associated signaling molecules. as well as provide a concise representation of the receptor life cycle, including internalization, degradation, and recycling processes.

The results of our modeling and simulation study reveal that canonical Wnt signaling is not characterized by one, but several internalization mechanisms, which all play varying yet pivotal roles in early, mid-term, and long-term regulation. Our results suggest a previously underestimated role of receptor trafficking and recycling, particularly during mid-term and long-term regulation. The observation that receptor distribution, in general, is much sparser than the membrane compartmentalization by actin-cytoskeleton implies that receptor complexes have to overcome several membrane fences to co-localize. Therefore, effective receptor clustering relies on directed trafficking via internalization and fast recycling particularly to form larger protein complexes, as suggested by our simulation results.

To better understand our simulation results, we analyzed the firing rate of each individual rule defined in our simulation model throughout the simulation runs. In Figure 10, the main processes with respect to receptor trafficking implemented in our model are illustrated with arrow color (gray and red) referring to the simulation models and arrow width being scaled with the mean firing rate of the corresponding rule in our simulation model. Here, it becomes obvious that in a model without receptor trafficking (gray arows) the majority of trimeric Wnt/Fz/LRP6 receptor complexes (TRC) and small multimeric receptor complexes (MRC) are internalized before they can merge with each other. This limits the aggregation of TRC into larger MRC clusters, hence limits the size of MRC. Visualizing the rule count distribution of the model with receptor trafficking (red arrows) reveals a more even distribution of rule firing rates, indicating a more balanced and fine-tuned activation of various, concurrent mechanisms. Lateral association of TRC (dimerization) as well as intracellular transport (i.e., recycling of early endosomes) contribute to and promote the assembly of stable signalosomes. On the other hand, the intensity of the signal activation is limited and controlled by caveolin-mediated degradation of signalosomes as well as early ZNRF3 and RNF43 induced clathrin-dependent internalization and degradation of TRC (Fig. 10).

**Fig. 10:**
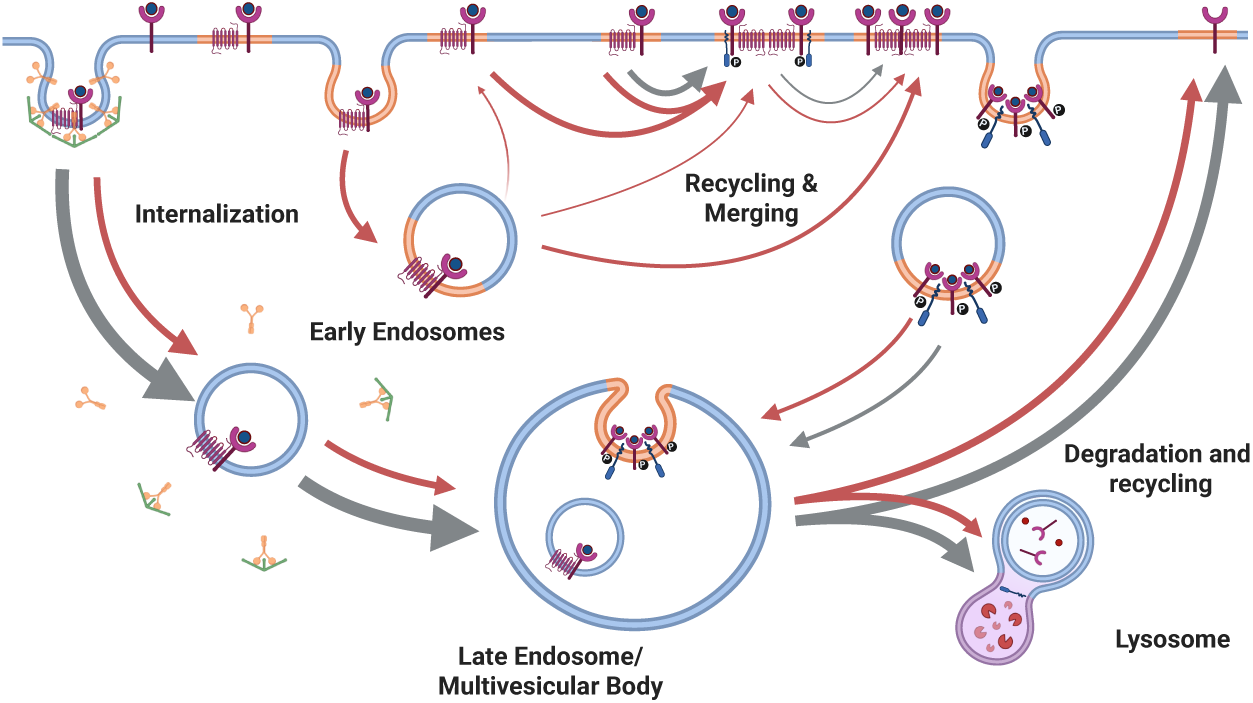
Comparison of rule counts of models with (red arrows) and without (gray arrows) intracellular receptor traffic. Arrow width scales with rules fired in simulations of the corresponding model (indicated by arrow color). Created in BioRender. Haack, F. (2025) https://BioRender.com/dymsp27

Lastly, fast receptor trafficking might also explain the primary localization of Wnt proteins in raft domains several hours after Wnt stimulation or in transfected, Wnt-producing cells, as observed in [68] and [69]. While Wnt-receptor complexes outside of raft domains are quickly internalized and degraded, raft-specific recycling and protein sorting steer the localization and enrichment of Wnt-receptor complexes to raft domains. This process results in the predominant localization of Wnt proteins and associated protein receptor complexes to raft domains. Once established, large signalosomes are eventually internalized, and important components of the destruction complex, which have been recruited to the membrane, are sequestered in multivesicular bodies and eventually primed for lysosomal degradation. However, without continuous rounds of receptor recycling, long-term Wnt signaling leads to the depletion of cell surface LRP6 receptors, which lowers receptor density and signal activation efficiency. Our model analysis thus reveals that raft-associated signalosome formation and elevated recycling lead to prolonged *β*-catenin signaling, while lowering receptor trafficking and recycling dynamics effectively attenuates pathway activation in terms of *β*-catenin aggregation.

While small receptor assemblies, ranging from dimers to single tetramers, can form independently of endocytic processes under low Wnt stimulation, the formation of large signalosomes and the establishment of stable, long-term *β*-catenin activity apparently require active endosomal sorting and recycling. Targeting these trafficking processes therefore offers a strategy to selectively dampen pathological signaling while preserving low-level pathway activity in healthy cells. Notably, internalization and receptor trafficking represent a defining feature of aberrant Wnt-independent signaling in certain APC-mutant cancer cells, as recent studies uncovered [35, 115]. Further, the membrane composition, particularly in terms of cholesterol and fatty acid (omega-3 and omega-6) content, significantly impacts signalosome formation and internalization [116]. In summary, our integrative study complements existing experimental work by providing a mechanistic framework that links membrane composition, receptor trafficking, signalosome organization, and pathway output; thus providing an answer to the long-standing question of why endocytosis is essential for signal activation and why receptor internalization can either promote or attenuate pathway activity depending on the localization of the signaling complex. By bridging molecular-scale dynamics with long-term signaling behavior, our approach enables a more predictive understanding of how Wnt signaling is selectively amplified in disease. As such, it provides a conceptual framework for the development of targeted interventions in Wnt-driven diseases.

## 4 Methods

Due to its central role in numerous biological processes and cellular regulations, as well as in its aberrant form, in cancer development, the Wnt signalling pathway has been the subject of various simulation models [45]. However, membrane dynamics have only been considered in a few models so far [46–50]; none of them comprises an extensive, pathway-specific representation of receptor trafficking in Wnt signaling. Modeling the receptor life cycle, including receptor/protein complex formation, ligand – or raft-induced internalization, recycling, and intracellular sorting, poses a major challenge for computational representations of canonical Wnt signaling. These processes, together with the resulting spatial heterogeneity in receptors, kinases, and other signaling components, critically shape the pathway dynamics. To capture these processes, we employed a compartmental modeling approach using ML-Rules, a multilevel rule-based language [51, 52].

### Rule-based modeling based on ML-Rules

To develop the model, a modeling approach was required that supports dynamic compartmentalization, including the creation of new compartments, their deletion, and merging. Incorporating compartmental dynamics poses significant challenges for both the design of modeling languages and the underlying simulation engines. Many simulation tools in cell biology offer static compartments that do not change. This allows the modeler to consider the causal influence of compartments on the processes within. However, to account for the causal influence of the occurring processes on the compartmental structure and interactions, dynamic compartments are needed. ML-Rules is a rule-based language for multi-level cell biological modeling and supports a wide range of compartmental dynamics [51]. Various simulators have been developed for ML-Rules over the last decade [52]. Here, we use a more recent ML-Rules implementation in Rust that offers the required efficiency to model 1000s of compartments [53]. For more details on the implementation of the simulator, as well as synthax and semantics definition of ML-Rules, the interested reader is referred to [53**?**].

In the present model cellular and subcellular structures, such as the plasma membrane, raft domains, endosomes, and lysosomes, are represented as nested compartments with variable molecular content and attributes (e.g., radius, volume). ML-Rules enables the dynamic creation, merging, modification, and removal of compartments during run-time, allowing for a straightforward representation of processes such as endocytosis, recycling, and lysosomal degradation through declarative reaction rules. Below is an exemplary rule, that encodes the internalization of a lipid raft compartment.

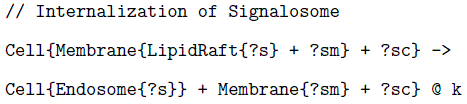

Note that compartments may be nested and may contain an arbitrary number of species or other compartments. In the upper rule the content of a *LipidRaft* compartment, located in the *Membrane* enters the *Cell* and its content (denoted as?s) is transferred to the newly (dynamically) created compartment *Endosome* with the reaction rate constant k. The remaining species of the other compartments cell and membrane remain unchanged. Reaction kinetics may depend on compartment attributes, enabling volume-dependent rate kinetics and dynamic redistribution of molecular species. This approach provided us with the means to mechanistically incorporate the essential steps of the LRP6 receptor life cycle into a coherent, quantitative model.

### Data sources and model integration

During the process of model development, we integrated experimental data and qualitative information from numerous studies (see Fig. 11). These data sources range from qualitative information to semi-quantitative, quantitative, or time-dependent measurements, mainly drawn from in-vitro experiments. The type of data source determined how information or data were integrated into the process of model development. While qualitative information could only be used to structurally shape the qualitative model, hence define the different entities and their connections (in terms of rule definitions), quantitative data could be used directly as parameter values (see Table 2) or, in the case of time-dependent measurements, as reference values for parameter value calibration. Thereby, careful considerations were required to integrate the data adequately, because numerous experimental variables can influence cellular responses. Differences in cell lines, culture conditions, and timing/dosing of treatments significantly alter outcomes; and variations in genetic manipulation methods, protein expression levels (endogenous, over-expressed), dosing strategies, and incubation durations further contribute to inconsistency among experimental measurements. To structure this large set of heterogeneous data we defined model requirements as part of a conceptual model that guided our model development [117]. These model requirements describe qualitative experimental observations or (semi-)quantitative measurements that should be reflected in the model. Typically these dynamics emerge during the simulation of the model, such as protein residence times at the cell surface, internalization dynamics or signalosome/lipid rafts size distribution. Table (1) describes a subset of model requirements, that were used during model development. These requirements were, for instance, used as reference points or parameter value boundaries during model calibration, or for face validation of certain model dynamics.

**Fig. 11:**
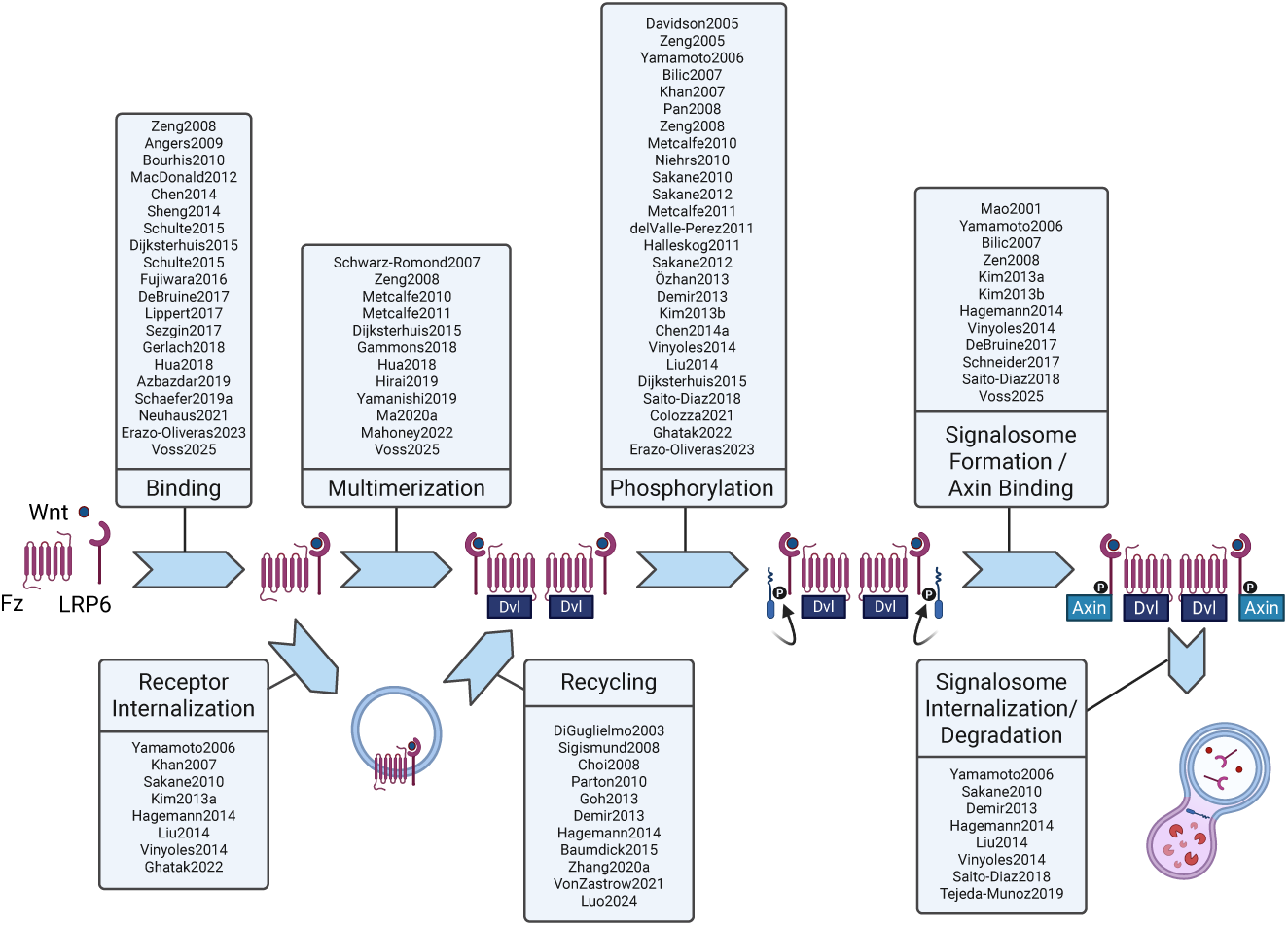
Graphic overview illustrating the integration of experimental data during model development. Arrows connecting model species represent submodels that describe corresponding mechanisms in the early steps of canonical WNT signaling, that are represented in the model, as described in the results section. Associated boxes list selections of references, from which either qualitative or semi-quantitative information was integrated to develop the respective part of the model or fit the corresponding kinetics.

### Model parameter calibration and validation

Direct measurements of kinetic or dissociation rate constants are most valuable for model development. In this case parameter values can typically be directly used as parameter values in the model. Table 2 lists all parameter values employed in our model. If experimental measurements exist or if parameter values were reused from previously published model components, particularly regarding intracellular down-stream signaling, the corresponding references are listed in the table. Parameter values that have been fitted in the course of the model development are marked with a star (*). The remaining parameter values for which experimental measurements or literature were unavailable, were fitted against various sources of experimental data using a genetic optimization algorithm that iteratively refined candidate parameter sets to reproduce data from in-vitro measurements. Starting off with a vector of initial candidate parameter values, i.e., the initial population, the simulation model was executed and its output was evaluated using a composite objective function. Note that our objective function consisted of multiple equally weighted distance metrics, each quantifying the deviation between simulated results and distinct experimental observations. This approach allowed us to simultaneously fit our model against various time-resolved in-vitro measurements on dimerization of trimeric receptor complex [95], LRP6 phosphorylation [71, 119, 120], and LRP6 internalization [32, 73, 74]. These metrics, i.e., the root mean square deviation of each model observable from the corresponding experimental data, were averaged across the available time points, normalized, and transformed into a fitness score that was used in the selection process of the population. Across successive generations, individuals with higher fitness were preferentially selected for crossover and mutation, generating new candidate solutions while maintaining population diversity. This evolutionary process continued until the convergence criteria (simulation trajectory of each observable lies within the measurement error bounds for all time points) were reached, at which point the best-performing parameter vector was identified as the calibrated model configuration.

After successfully calibrating the model parameter values related to receptor trafficking, we evaluated the predictive power of our model with regard to downstream signaling and pathway activation in terms of *β* −*catenin* stabilization and conducted a cross-validation experiment by mimicking the in-vitro experiments performed in [129]. We initialized our model with the same range of Wnt concentrations as applied in [129] and measured the accumulation of intracellular (total) *β*-catenin 2 hours after the start of the stimulation. As depicted in Figure 12, the response curve is in good agreement for all Wnt3a concentrations applied. Thus, our model, which has been fitted against experimental data retrieved from various experimental sources, is also capable of reproducing *β*-catenin kinetics reported for different Wnt3a stimuli. This is particularly noteworthy, as the model was calibrated exclusively using data from early signaling events, i.e. receptor complex dimerization, phosphorylation, and internalization, without incorporating downstream signal transduction data. Yet the model accurately captures downstream signal transduction and *β*-catenin stabilization.

**Fig. 12:**
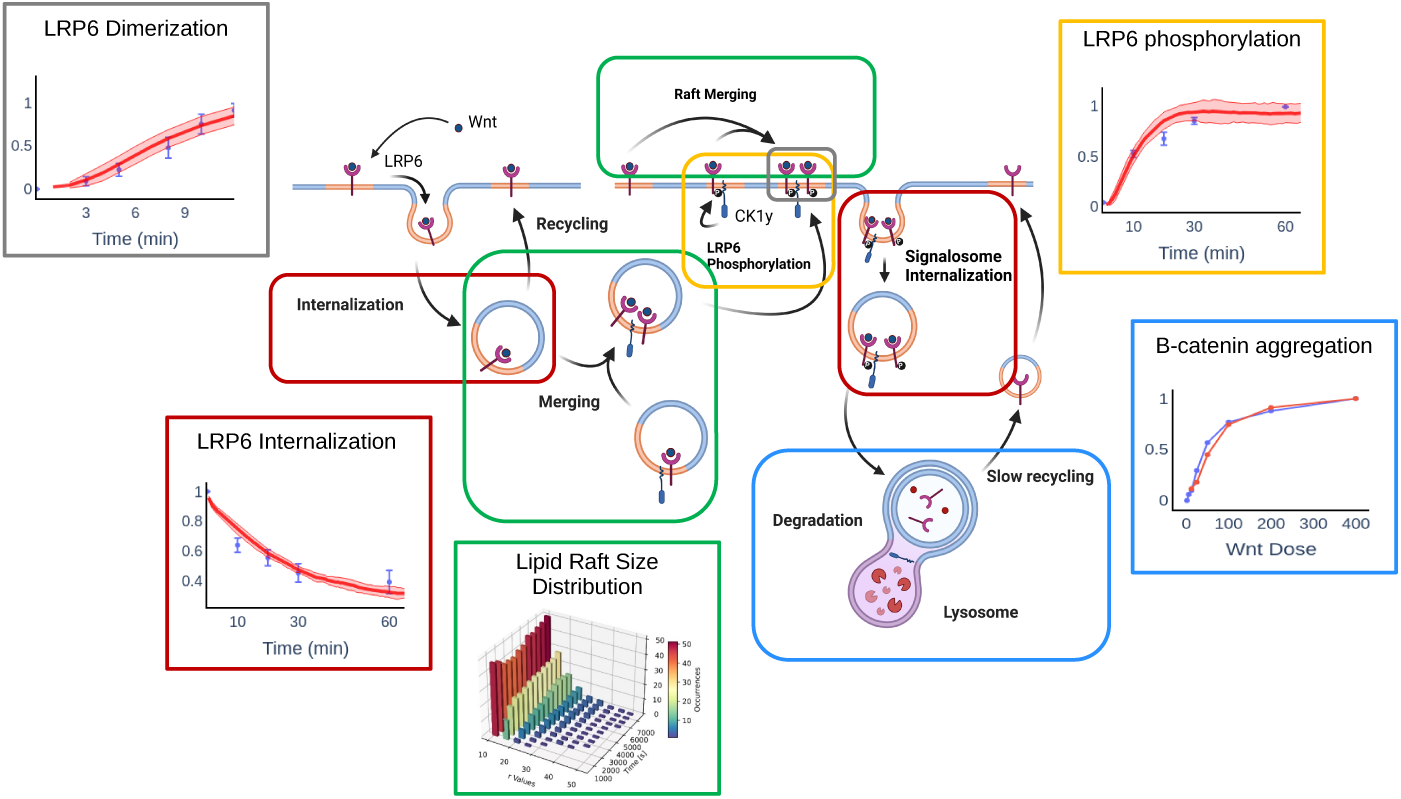
Model elements fitted against experimental data from various sources, including LRP6 phosphorylation (yellow), receptor dimerization and signalosome formation (green) and LRP6 internalization. In-vitro data of *β*-catenin aggregation obtained 2 hours after onset of Wnt stimulus for varying Wnt concentrations was used to validate the calibrated model. In the individual graphs, red lines correspond to simulation trajectories, whereas blue points or blue lines represent experimental measurements. Created in BioRender. Haack, F. (2025) https://BioRender.com/cwdveej

## Supplementary information

Supplementary information can be provided on request.

## Acknowledgements

The authors thank Lena Cibulski and Stefan Bruckner for their excellent technical support on visualization and analysis of time-dependent model observables using parallel coordinates. K.B. would like to acknowledge the Alexander von Humboldt Foundation for its awarding of the Humboldt Forschungspreis to work with Professor A.M.U.

## Declarations

### Author contribution

Conceptualization: FH, AMU, KB; Methodology: FH, AMU, KB; Data Curation: FH; Model Development and and Simulation Experiments: FH; Validation: FH; Visualization: FH; Writing – Original Draft: FH, AMU, KB

### Data availability

See Code availability.

### Code availability

Source code of the model and all relevant submodels as well as computational framework to execute the model simulations and analyse simulation results are provided with this submission to reproduce simulation results of all relevant figures. Upon publication all source code will be deposited on a public repository using permanent identifiers. Currently the code is available in the following git repository: https://git.informatik.uni-rostock.de/mosi/wntreceptortraffic

